# Climate and predation drive variation of diel activity patterns in chacma baboons (*Papio ursinus*) across southern Africa

**DOI:** 10.1101/2024.11.05.622068

**Authors:** L. Dzingwena, L. Thel, M. Choisy, R. Garbett, A. Wilkinson, J. Venter, H. Fritz, E. Huchard, F. Prugnolle, V. Rougeron

## Abstract

Understanding how animals adjust daily activity to environmental gradients reveals key drivers of behavioral plasticity. While diel activity is theorized to reflect trade-offs among thermoregulation, energy balance, and predation risk, few studies test these interactions at broad spatial scales within species. We investigated this in chacma baboons (*Papio ursinus*) using over a million camera-trap detections across 29 sites in six biomes (2016–2022) in South Africa and Zimbabwe. Activity declined by about 3% with latitude, consistent with lower resource predictability and increased with thermal stress, while sleep and wake times were similar across sites. Baboons avoided midday heat but increased dawn and night activity under predator pressure. These findings show how abiotic and biotic pressures shape diel schedules and highlight temporal flexibility as an adaptive strategy for generalist mammals in a changing world.

## Introduction

Wildlife’s diel activity patterns, referring to the timing and vigor of activity throughout a 24-hour cycle, vary latitudinally among and within species (1,2). These patterns are shaped by both intrinsic factors such as circadian rhythms (3,4), and extrinsic factors such as environmental conditions (5,6) and inter-species interactions (7,8). Although activity throughout the year is shaped by large-scale phenomena, like the spatio-temporal accessibility of food resources (9) or reproductive timing (10), daily activity is fine-tuned to meet energy requirements (3), avoid thermal stress (10), minimize actual (7) or perceived (11) predation risk, and mitigate competition (12). Diurnal activity allows better predator detectability, thermoregulation through sun exposure or shade seeking, and habitat exploration, while nocturnal activity provides benefits such as cooler temperatures and reduced competition (13). These daily activity strategies, often flexible within individuals and populations, reflect not only behavioral plasticity but also underlying phenotypic adaptations, such as differences in visual acuity, thermoregulatory capacity, or metabolic rates, evolved to optimize performance in specific ecological contexts (6,14,15).

Mammalian diel activity patterns reflect adaptive trade-offs among energy acquisition, thermoregulation, and risk avoidance, shaped by environmental factors like climate, habitat structure, and predator assemblages (11,13,16). For instance, desert ungulates such as gemsbok (*Oryx gazella*) and Arabian sand gazelle (*Gazella marica*) limit activity to cooler periods to avoid extreme heat, despite increased nocturnal predation risk (17,18). Conversely, species in temperate or complex habitats, like elephants (*Loxodonta africana*) and red deer (*Cervus elaphus*), maintain broader activity windows to optimize foraging and thermoregulation (19,20). Predator-prey dynamics further influence activity patterns; prey like impalas (*Aepyceros melampus*) and warthogs (*Phacochoerus africanus*) adjust their activity to reduce overlap with nocturnal predators such as lions (*Panthera leo*) and spotted hyenas (*Crocuta crocuta*) (21,22). In densely vegetated areas, ambush predators like leopards (*Panthera pardus*) affect prey activity (22), while predators like African wild dogs (*Lycaon pictus*) and caracals (*Caracal caracal*) may shift their own activity to align with prey availability (23,24). In stable environments with lower predator densities or better habitat visibility, animals may experience reduced predation pressure, allowing for more extended or diversified activity without substantially increasing vulnerability (23,24). Thus, diel activity patterns emerge from complex ecological and evolutionary interactions, rarely dictated by a single factor. While thermoregulation and food availability may be the strongest constraints in arid or resource-scarce systems, risk avoidance may take precedence in predator-dense areas

This complexity is also observed among non-human primates (NHPs), whose activity patterns reflect similar trade-offs. Most anthropoid species (e.g. chimpanzees) are diurnal, while prosimians (e.g. lemurs, lorises, and tarsiers) exhibit a broader range of activity patterns, being either nocturnal, cathemeral or diurnal (25,26). Activity patterns in NHPs vary in response to predator-prey dynamics (27,28), time budgets constraints (29), and seasonal shifts in food availability or environmental cues like temperature and lunar cycles (30). For example, the cathemeral mongoose lemur (*Eulemur mongoz)* becomes more nocturnal in the dry season to manage heat stress and predation risk (31), likewise the owl monkey in Argentina shifts patterns seasonally (32). However, most studies focus on single populations, limiting our understanding of how broad-ranging generalist species adjust their daily activity patterns across different ecological environments.

Largest among baboons, highly adaptable and widely spread across diverse habitats ranging from savannas and grasslands to forests, deserts, and mountains (Fig. 1), the chacma baboon (*P. ursinus*) offers an interesting model for studying daily activity patterns across multiple environments (32–34). This species demonstrates marked behavioral flexibility optimizing active hours by balancing foraging, resting, grooming, and socializing, while avoiding predation, heat stress, and energy loss (35–37). For instance, in South Africa radio telemetry studies in the Soutpansberg Mountains found evidence of nocturnal activity in chacma baboons as a response to seasonal fluctuations in day length and lunar luminosity (36), while chacma baboons at DeHoop Nature Reserve rest and seek shade to avoid midday heat (38). During dry summers, they increase daily activity to compensate for food scarcity (35,39). Predators such as lions, leopards, and spotted hyenas, which are among the leading causes of non-anthropogenic adult baboon mortality (40,41), also influence chacma baboon spatio-temporal habitat use. For example, olive baboons delay morning activity to avoid leopards in Kenya (42). Therefore, chacma baboons provide an interesting model to study how environmental and ecological factors shape diel activity patterns, informing behavioral ecology and climate adaptation.

**Fig. 1.**
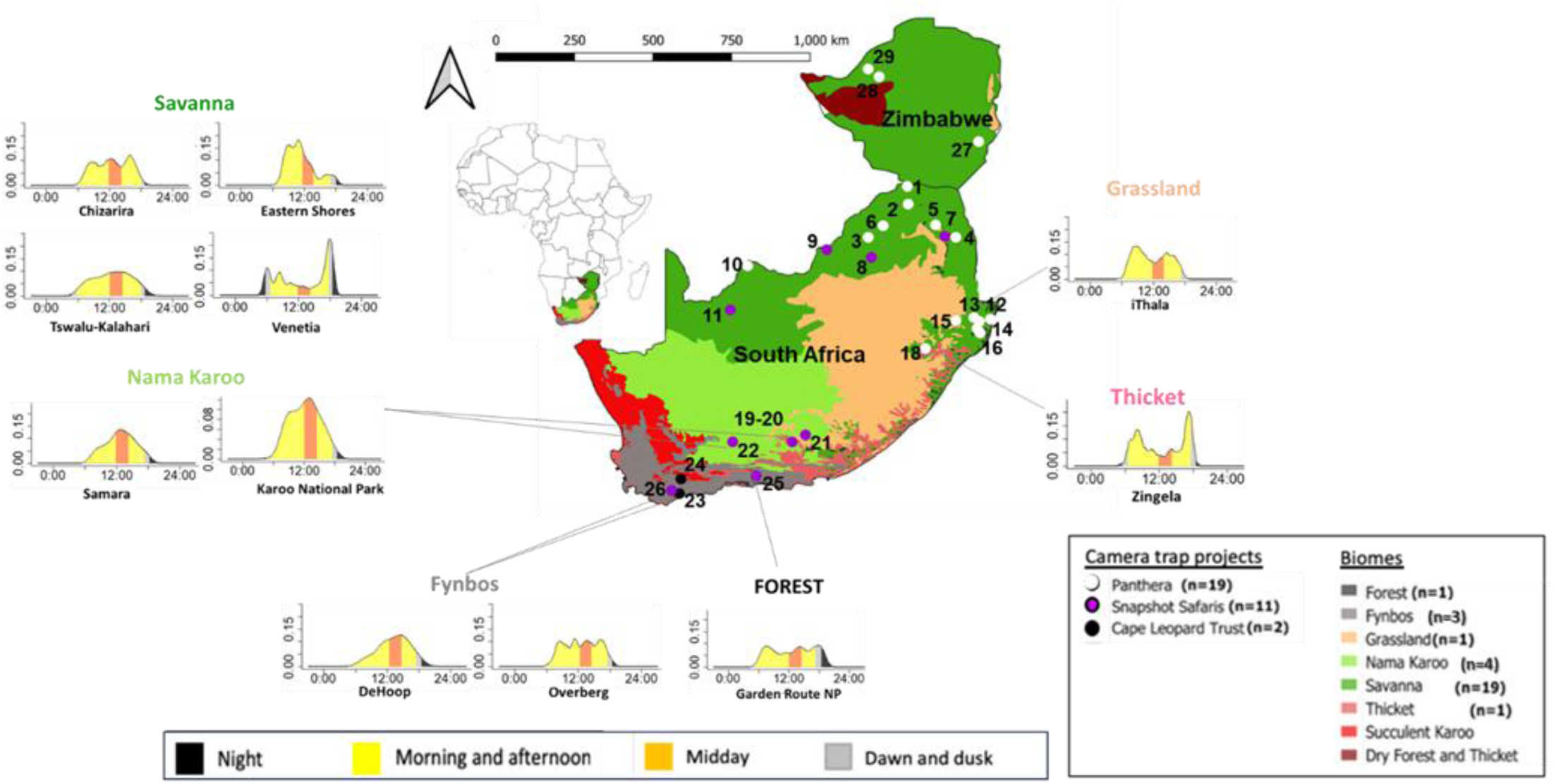
Examples of activity distributions of chacma baboons in different biomes and geographic distributions. The sources of camera trap data are indicated in white for Panthera, purple for Snapshot Safari and black for Cape Leopard Trust. The numbers on the map correspond to: 1. Venetia Limpopo Nature Reserve; 2. Lajuma Nature Reserve; 3. Welgevonden Game Reserve; 4. Timbavati Private Nature Reserve; 5. Makalali Game Reserve; 6. Lapalala Wilderness; 7. Associate Private Nature Reserves; 8. Pilanesberg National Park; 9. Madikwe Game Reserve; 10. Khamab Kalahari Reserve; 11. Tswalu Kalahari Reserve; 12. uMkhuze Game Reserve; 13. Phinda Private Game Reserve; 14. Manyoni Private Game Reserve; 15. Ithala Game Reserve; 16. Hluhluwe Game Reserve; 17. Eastern Shores; 18. Zingela Nature Reserve; 19-20. Samara Private Nature Reserve & De Nek Farm; 21. Mountain Zebra National Park; 22. Karoo National Park; 23. Overberg; 24. Little Karoo; 25. Garden Route National Park; 26. DeHoop Nature Reserve; 27. Save Valley Conservancy; 28. Chirisa Safari Area; 29. Chizarira National Park. The Y-axis represents the density of activity, and the X-axis represents the hours throughout the day (in solar time, see text for details). Each day is divided into six periods: dawn and dusk (grey), morning and afternoon (yellow), mid-day (orange), and night (black).

The aim of this study was to characterize diel activity patterns of chacma baboons across 29 sites spanning six distinct biomes (savanna, nama-karoo, fynbos, forest, thicket, and grassland) in South Africa and Zimbabwe, covering a latitudinal range from 34.46°S to 17.70°S. These sites differ markedly in both abiotic (e.g., temperature, precipitation, NDVI) and biotic (e.g., predator species) characteristics. This diversity provided a unique opportunity to investigate how chacma baboons adjust their activity patterns in response to local selective pressures related to thermoregulation, foraging efficiency, and predation risk. Utilizing an extensive camera trap dataset, we tested the following predictions: *Geographical perspective* H1) Chacma baboon diel activity levels vary geographically due to environmental differences. Specifically, we expect chacma baboons in southern regions, where climates are milder, more stable, and generally more productive year-round, to sustain higher activity levels throughout the day. In contrast, chacma baboons in northern regions, where environments are harsher, are expected to show lower overall activity levels, reflecting a strategy of energy conservation in response to elevated thermal stress and limited food availability. *Seasonal perspective* H2) Within sites, chacma baboon diel activity shifts seasonally. In summer, chacma baboons are expected to reduce activity during the hottest parts of the day (midday and afternoon) to avoid heat stress. In winter, when days are shorter and colder and food is less available, we expect chacma baboons to extend their active periods during daytime to increase foraging time and meet higher energy demands for thermoregulation. We then evaluated the contribution of specific environmental drivers (temperature, precipitation, NDVI and predator species’ activity patterns), by testing the following hypotheses: *Thermoregulation constraints* H3) Within sites, the timing of activity is expected to be influenced by temperature fluctuations throughout the day. In warmer environments, chacma baboons are expected to be more active during cooler day periods to avoid heat stress, while those in cooler environments remain active throughout the day. *Foraging optimization* H4) In environments with limited food resources, chacma baboons are expected to adjust their activity patterns by increasing movement or foraging effort during specific periods of the day when conditions are most favorable. These shifts may not lead to higher overall activity levels but rather reflect a strategic redistribution of activity during certain day periods to optimize energy intake under resource-scarce conditions. *Predation risk avoidance* H5) We hypothesize that chacma baboons adjust their activity patterns based on predator presence at specific day periods. Depending on the predator species, we expect: (i) leopards, as the main predator of chacma baboons, to exert the strongest influence, leading to reduced activity during dawn and dusk and nocturnal periods; (ii) spotted hyenas, being primarily nocturnal and opportunistic, to promote chacma baboon’s activity during day; (iii) lions, though not frequent predators of baboons, to indirectly reduce activity level at dawn and dusk through disturbance or perceived risk.

## Materials and method

### Study sites

Chacma baboons have a broad distribution across southern Africa, inhabiting countries such as South Africa, Namibia, Botswana, Zimbabwe, Mozambique, Zambia, and Angola. In our study, we utilized existing camera trap data from 29 sites across six distinct biomes in South Africa and Zimbabwe. Most of these sites (n=19) were situated within the savanna biome, while the remaining sites were distributed among the nama-karoo (n=4), fynbos (n=3), afro-temperate forest (n=1), grassland (n=1), and thicket (n=1) biomes.

The six biomes in this study; savanna, nama-karoo, fynbos, afro-temperate forest, grassland, and thicket, exhibit distinct environmental conditions influencing resource availability (see Table S1). Savannas feature seasonal rainfall (235–1,000 mm/year) and high temperatures, supporting grassy plains with scattered trees (43,44). The semi-arid Nama-karoo experiences low, variable rainfall (100–520 mm/year) and extreme temperature fluctuations (45,46). Fynbos, within a Mediterranean climate zone, has winter rainfall and dry summers, characterized by dense shrub cover (47). Afro-temperate forests maintain stable climates with moderate temperatures and higher precipitation (500–1,200 mm/year), supporting dense vegetation (44). Grasslands, dominated by grass with few trees, experience high precipitation seasonality and frequent fires. Thickets consist of dense shrubs and trees, occurring in areas with moderate rainfall (200–950 mm/year) (48). This environmental diversity provides a unique opportunity to examine how chacma baboons adjust their activity patterns in response to local selective pressures related to thermoregulation, foraging efficiency, and predation risk.

### Data collection

#### Camera trap data

In this study, we capitalized on pre-existing camera-trap sets deployed in Southern African regions for three projects, each with distinct objectives: Snapshot Safari, Panthera, and Cape Leopard Trust (*49–51*, Supplementary information). Depending on the site, cameras were active between 2016 and 2022, providing temporal coverage across multiple years and seasons (Table S2), with an average of 40 (minimum=9; maximum=77) camera stations per site (Table S2). Monitoring efforts spanned a median of 349 (minimum=124, maximum=1,681) days per site (Table S2). From this dataset, we extracted images of chacma baboons and of their primary predators, namely lions, leopards, and spotted hyenas.

#### Abiotic variables

To understand the ecological and environmental factors influencing chacma baboon’s diel activity patterns, we focused on three primary abiotic variables: (i) temperature, which is linked to heat stress, metabolic rates, and energy expenditure in animals (2,18); (ii) precipitation, which impacts water and food availability and distribution, and overall habitat quality (36,52); (iii) Normalized Difference Vegetation Index (NDVI), a proxy of primary productivity and food resource availability as well as habitat suitability (53,54). We extracted the average NDVI at the study site scale for 16-day periods from MODI13Q1.061 Terra Vegetation Indices 16_Day Global 250m dataset (https://lpdaac.usgs.gov) using Google Earth Engine (https://earthengine.google.com). Mean monthly rainfall was obtained from TerraClimate Monthly Climate and Climatic Water Balance for Global Terrestrial Surfaces dataset (55), and mean hourly temperatures were extracted from Era5-Land Hourly-ECMWF Climate Reanalysis dataset (56). We calculated the mean seasonal hourly temperature, mean seasonal NDVI, and mean seasonal rainfall for each study site, using a 500m radius buffer from the outermost camera trap stations positioned in each site. Seasonal temperatures, annual precipitation, rainfall seasonality and NDVI values for each site are summarized in Table S1.

### Camera trap dataset merging and filtration steps

A total of 1,771,424 camera trap images containing mammal and bird species (Table S2) were obtained from all 29 sites. This dataset was cleaned and filtered to remove duplicate records, correct date and time and exclude records with inconsistent date-time stamps (Fig. S1). Since recommended sample sizes for diel activity patterns are between 100 and 500 observations (57,58), we selected sites where at least 200 chacma baboon capture events were obtained and ended up with a final dataset comprising 29 out of the 61 sites initially considered (Figure 1, Table S2). This process resulted in a refined dataset of 1,460,847 capture events, of which 268,121 were chacma baboons, 11,849 were leopards, 8,999 were lions and 15,200 were spotted hyenas (Table S2). Predator presence at each site was determined based on filtered capture events from camera traps, identifying which predator species (lion, leopard, spotted hyena) were detected at each location. Only sites with at least 100 capture events of a certain predator were then considered for the analyses of the effect of this predator’s activity on chacma baboon activity (n=25) (57).

### Chacma baboon diel activity patterns modelling

Activity was defined as any event where at least one individual was detected by a camera trap. The timestamp of the first image per event was used. To adjust for latitude and seasonal variation, times were converted to solar time using the ‘solartime’ function in the R package *activity* (59), standardizing day length across sites (9.8–14.5 hours). Following Peral, Landman and Kerley (2022), no temporal filters were applied, preserving full temporal resolution for accurate activity analysis(60).

To characterize diel activity patterns in each studied site, non-parametric kernel density functions were applied using the ‘fitact’ function from the R package *activity* (59), estimating activity levels as the proportion of time baboons were active during a 24-hour period, with ***Activity level = 2πfmax***, where ***2π*** represents time in radians and ***fmax*** is the maximum of the kernel density plot of chacma baboons per site. Diel activity distributions were assessed at each site using the ‘densityPlot’ function in the R package *overlap* (61). The resulting ‘activity distribution’ describes the kernel density curve shape over a 24-hour period.

To further analyze chacma baboons’ diel activity patterns, six parameters were extracted: activity level estimate, activity distribution, time and height of maximum peak of activity, wake up and sleeping time. Using the ‘findpeaks’ function from the R package *pracma* (62), we identified the timing and amplitude of activity peaks. Activity level estimate was defined as the proportion of time animals were active over the 24-hour cycle, calculated as the area under the kernel density curve of capture times (58). “Wake-up” time was defined as the first inflection point, and “sleeping time” as the last deflection point, determined by calculating the first and second derivatives of the kernel density curve.

Given seasonal variation in chacma baboon activity (63,64), capture events were grouped into four seasons: autumn (Mar–May), winter (Jun–Aug), spring (Sep–Nov), and summer (Dec–Feb). Seasonal activity levels per site were extracted using kernel density functions from the R package *activity* and combined unless stated otherwise.

To refine activity estimates, the day was divided into six periods: (i) dawn (1 hr. before sunrise to sunrise), (ii) morning (sunrise–noon), (iii) midday (noon–2 hrs. after), (iv) afternoon (post-midday to dusk), (v) dusk (sunset–1 hr. after), and (vi) night (1 hr. after sunset to 1 hr. before sunrise) (7,65). Sunrise, noon, and sunset times for each event were calculated using location and date via the ‘getSunlightTimes’ function in the R package *suncalc* (66). Noon was defined as the time when the sun reaches its highest point in the sky (local celestial meridian). To obtain fractions of activity level estimates per each day period in each season across sites, we used the function ‘integral’ in R package *pracma*, which computes the numerical integration 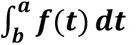, where ***f***(***t***) is the activity kernel density function and ***a*** − ***b*** is the corresponding time interval for each day period.

### Statistical analysis

#### Chacma baboon’s diel activity patterns across sites and seasons (H1-2)

To assess whether chacma baboon activity patterns vary geographically (H1), we analyzed the influence of latitude on five parameters: activity level estimate, timing and magnitude of peak activity, wake-up time, and sleep time. We employed beta regression models for activity level estimates and linear regression models for the other parameters. Model assumptions, including normality and homoscedasticity of residuals, were evaluated using the ‘simulateResiduals’ function from the R package *DHARMa* (67). Beta regression models, implemented via the *glmmTMB* package (68) were used to examine the relationship between total activity level estimates and latitude. Additionally, we investigated seasonal variations (summer, autumn, winter, spring) in activity level estimates using beta regression models. To explore differences in activity levels among sites during specific periods of the day (dawn, morning, midday, afternoon, dusk, night), we applied the Kruskal-Wallis’s test. Then, we used beta regression models to assess seasonal variations in activity levels within each diel period, followed by Tukey’s post-hoc tests to identify pairwise seasonal differences. Due to the uneven distribution of sampled sites across biomes (ranging from 1 to 19 sites per biome; see Fig. 1), we did not perform comparative analyses of activity patterns among biomes.

#### Variables influencing chacma baboon’s diel activities (H3-5)

Firstly, we tested the additive effect of three abiotic factors (temperature, precipitation and NDVI) on the activity level estimates of chacma baboons using beta regression models (’glmmTMB’ R package; (68)). We checked the preliminary assumptions of beta distribution and homoscedasticity of the residuals of the model using the function ‘simulateResiduals’ in the R package *DHARMa* (67). We checked the absence of multicollinearity among the explanatory variables with the variance inflation factor using the ‘check_collinearity’ function in the R package *performance* (69). The best model was selected for interpretation after removing outliers (maximum of 2 outliers per period) from all time periods except dawn and night, where no outliers were detected. To ensure comparison, all explanatory variables were standardised by subtracting their means and dividing by the global standard deviation.

Separate beta regression models were run for each predator species (lion, leopard, spotted hyena) to test how their seasonal and period-specific activity influenced baboon activity, with abiotic covariates included as additive effects. Model assumptions and multicollinearity were checked as above. Analyses focused on sites where at least one predator was detected. Predator activity levels were estimated using the same method as for baboons.

Finally, we quantified the temporal tolerance and avoidance of chacma baboons toward predators by examining the overlap coefficients, a unitless value ranging from 0 (no overlap) to 1 (complete overlap). To compare the probability density functions between chacma baboons and each predator species in each site, and test if they arise from the same distribution (i.e. are characterized by a significant overlap), we used the function ‘CompareCkern’ in the R package *activity* (59). We used bootstrapping (n=500 iterations) to obtain 95% confidence intervals (*65*). We plotted the activity curves for chacma baboons and each predator across all sites using the ‘densityPlot’ function in the *overlap* R package (61) (Fig. S3). All statistical analyses were done in R software version 4.3.1 (70).

## Results

### Chacma baboons are more active in the mornings and afternoons

Our analysis revealed that chacma baboons’ activity level estimates vary among day periods (dawn, morning, mid-day, afternoon, dusk, and night) within a 24-hour day cycle. We found that chacma baboons were most active (52.5% of all detections) and showed most variation across sites (range=8-84% of the day during which they were active; Fig. 2) in the morning, and then during the afternoon (20.8% of all detections), followed by mid-day (17.6% of all observations), dawn (5.8%), night (2.9%) and eventually dusk (0.3% of all detections; Fig. 3).

**Fig. 2.**
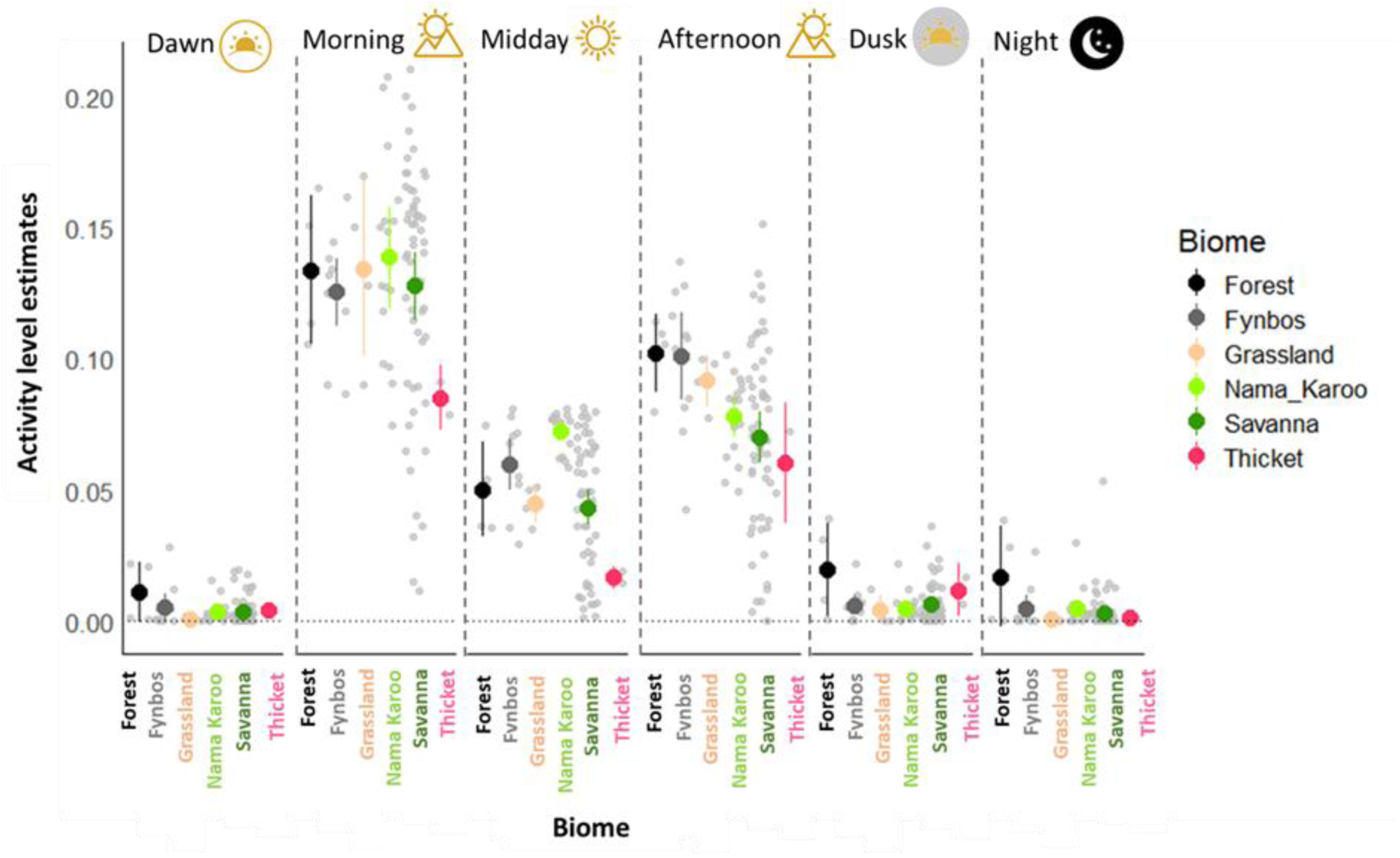
The biome specific average diel activity level estimate of chacma baboons per day period (i.e. dawn, morning, mid-day, afternoon, dusk and night). In each caption, light grey dots show site-specific activity levels. Colored dots represent biome-level averages: Fynbos (grey), Forest (black), Nama-karoo (light green), Thicket (pink), Grassland (peach), and Savanna (green). Vertical lines show 95% confidence intervals; wider intervals reflect greater within-biome variation.

**Fig. 3.**
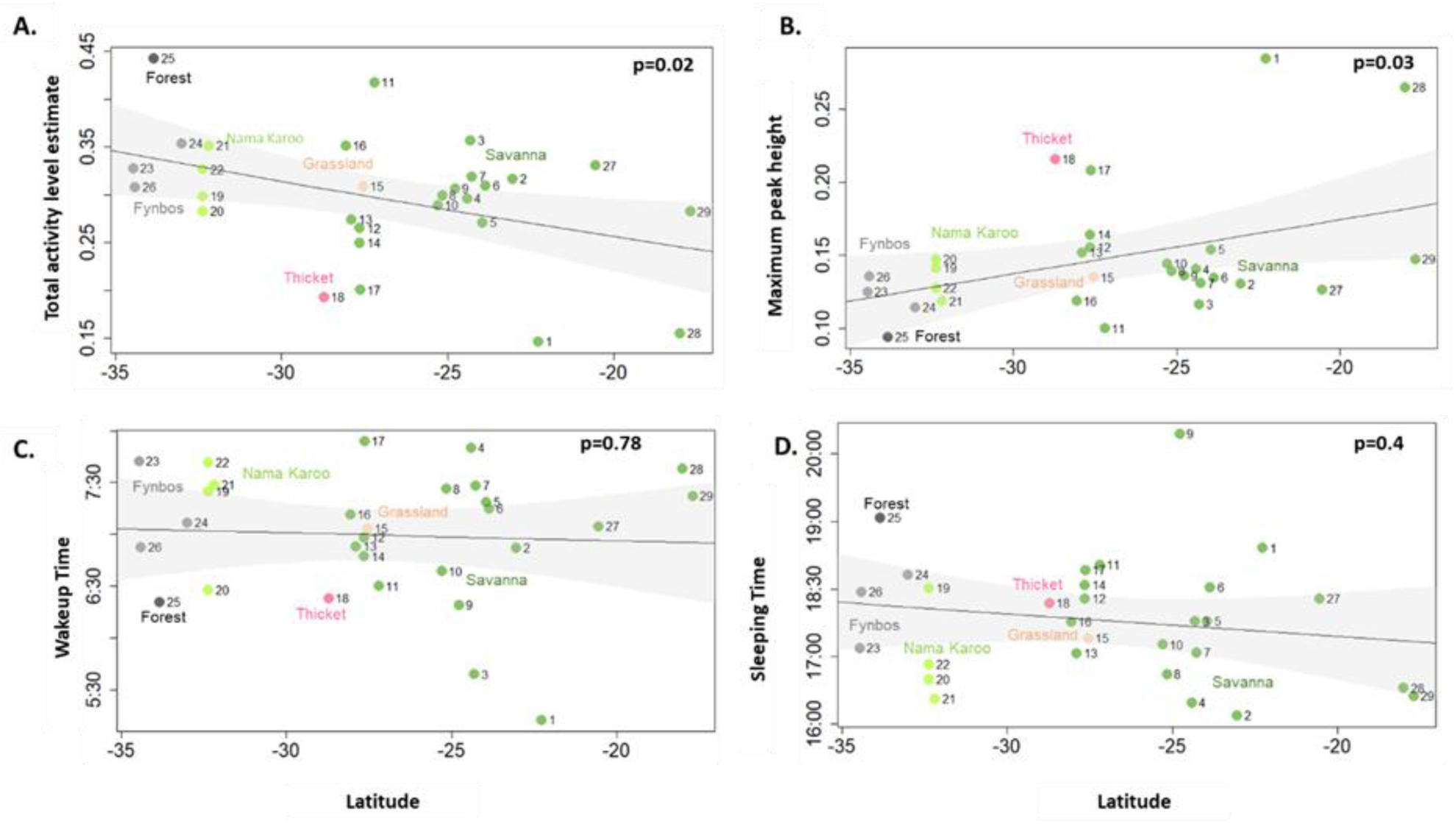
Diel activity patterns of chacma baboons of the 29 study sites across a latitudinal gradient and six Southern African biomes: Fynbos (grey), Forest (black), Nama-karoo (light green), Thicket (pink), Grassland (light peach) and Savanna (green). **(A)**. Total activity level estimates. **(B)**. Height of maximum peak of activity. **(C)**. Wake up time. **(D)**. Sleeping time. The dots correspond to the raw data, the solid line represents the predicted values from the best model, the shaded area represents the 95% confidence intervals estimated from the model.

### Differential diel activity patterns across sites (H1)

Chacma baboon activity level estimates significantly varied among sites for almost every period of the day, namely dawn, morning, mid-day and night (Table 1). In the afternoon this variation was only marginally non-significant (Table 1). No chacma baboons’ activity was detected at night in Samara, De Hoop, Chirisa, Chizarira and Associate Private Nature Reserves. Similarly, there were no chacma baboon detections in De Hoop, Chizarira and Samara at dawn, and in more than half of the savanna sites at dusk.

**Table 1:**
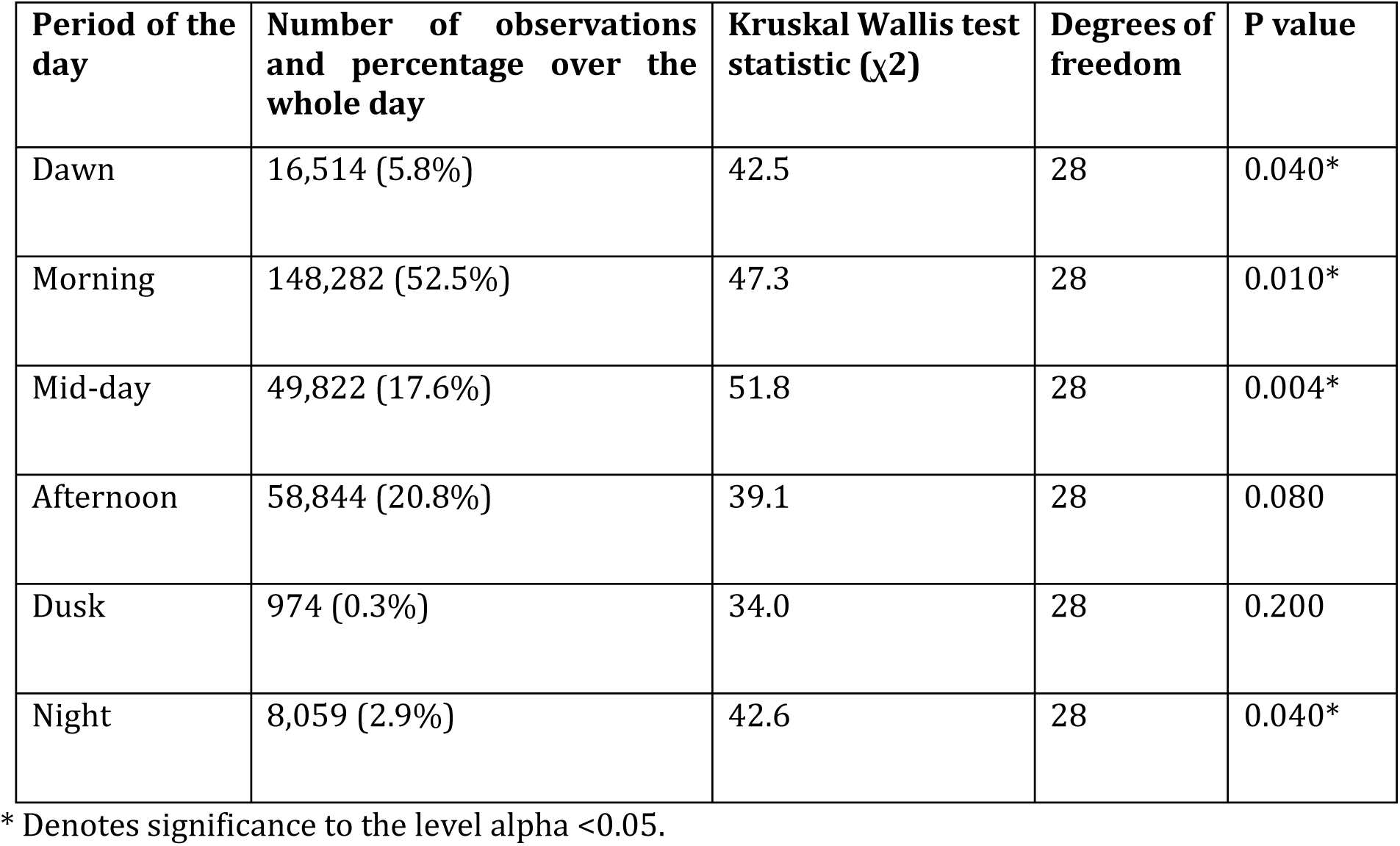
Difference in activity level estimates among sites in different periods of the day (Kruskal Wallis test).

| Period of the day | Number of observations and percentage over the whole day | Kruskal Wallis test statistic ($\chi^2$) | Degrees of freedom | P value |
| --- | --- | --- | --- | --- |
| Dawn | 16,514 (5.8%) | 42.5 | 28 | 0.040* |
| Morning | 148,282 (52.5%) | 47.3 | 28 | 0.010* |
| Mid-day | 49,822 (17.6%) | 51.8 | 28 | 0.004* |
| Afternoon | 58,844 (20.8%) | 39.1 | 28 | 0.080 |
| Dusk | 974 (0.3%) | 34.0 | 28 | 0.200 |
| Night | 8,059 (2.9%) | 42.6 | 28 | 0.040* |

\* Denotes significance to the level $\alpha < 0.05$ .

Among the 29 sites, chacma baboons showed the highest total activity in Garden Route National Park (0.44±0.01, forest biome) and the lowest in Venetia (0.15±0.005, savanna biome) (Fig. 1, Fig. 3A). Peak activity heights varied across sites within biomes (Fig. 1, Fig. S2), with higher total activity in southern than northern regions (Fig. 3A). Activity decreased by 2.84% per degree of latitude from south to north (β=0.028±0.01, p=0.02).

Trimodal activity was observed in 9 of 19 savanna sites and Garden Route, with morning peaks (0530–1127 hrs.) in savannas and dusk (1815 hrs.) in the forest biome (Fig. 3B). Unimodal patterns occurred in Tswalu (afternoon, 1428 hrs.), Madikwe (mid-day, 1303 hrs.), and Associate Reserves (morning, 0931 hrs.), as well as all nama-karoo and in De Hoop (fynbos, 1159 hrs.). Bimodal activity appeared in iThala (0820 hrs., grassland) and Zingela (1718 hrs., thicket) (Fig. 1). In the fynbos, baboons showed multimodal activity: Overberg (1124 hrs.) and Little Karoo (1539 hrs.) (Fig. 1). Peak height increased with latitude (β=0.004, p=0.03; Fig. 2B), indicating stronger activity peaks in northern sites.

Wake-up and sleep times were unaffected by latitude after adjusting for sunrise/sunset (wake-up: β=-0.01, p=0.78; sleep: β=-0.03, p=0.4; Figs. 2C, 3C, 3D).

### Seasonal differences in the timing of chacma baboons’ activity (H2)

While total daily activity level estimates did not differ significantly among seasons (summer, autumn, winter, spring; p = 0.76), results showed significant seasonal differences in chacma baboons’ activity when analyzed by specific day periods, except for mornings (p=0.18, Fig. 4) and mid-day periods (p = 0.34, Fig. 4). Specifically, chacma baboon activity at dawn, dusk, and night was significantly higher in winter and autumn compared to summer and spring (Fig. 4). By contrast, afternoon activity levels showed mixed patterns: while there were no significant differences between autumn and other seasons, summer showed significantly lower afternoon activity than spring and winter (Fig. 4, Table 2).

**Fig. 4.**
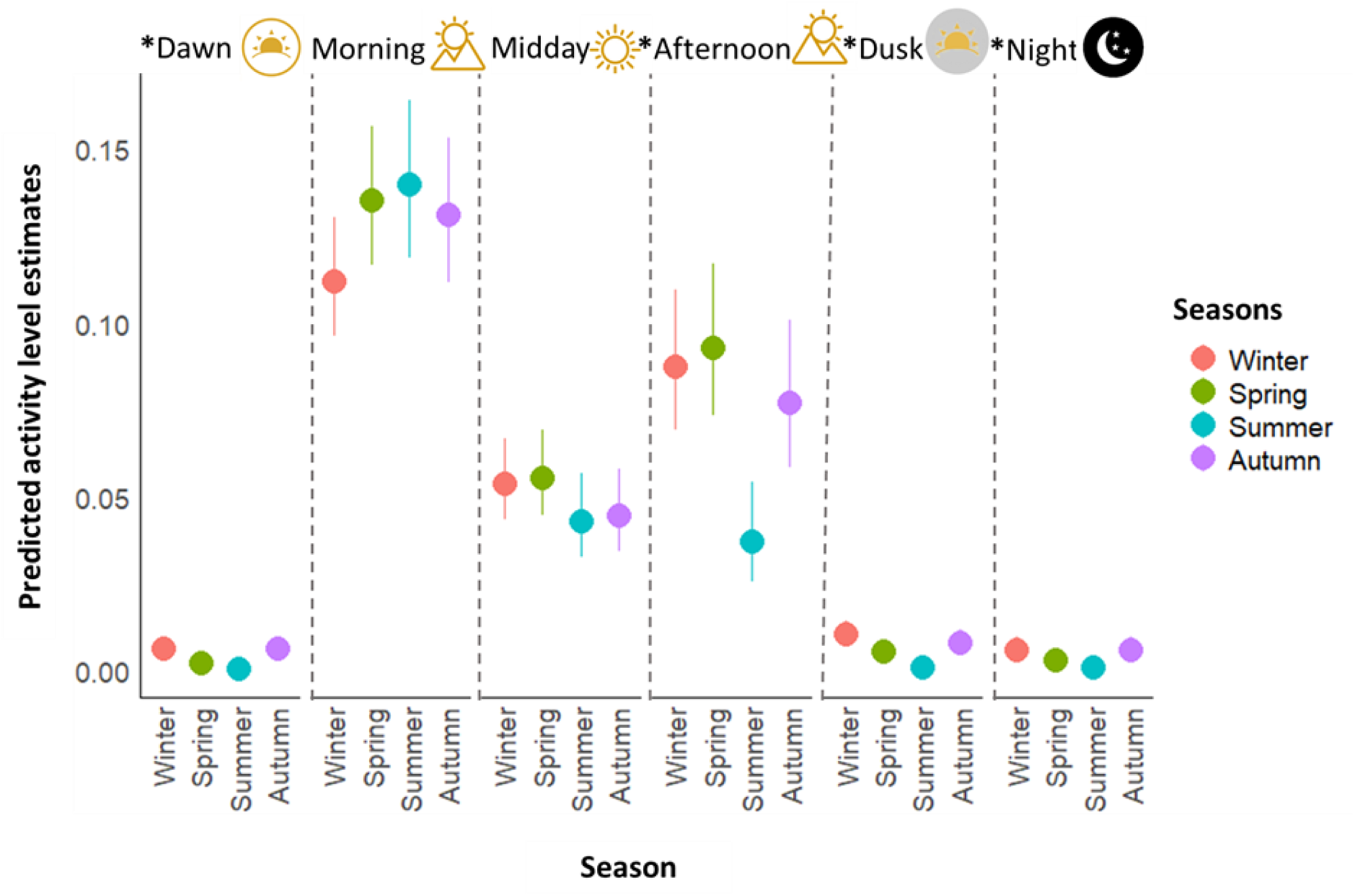
Seasonal variation in the total activity level estimates of chacma baboons in each day period (i.e. dawn, morning, midday, afternoon, dusk and night): Within each caption, colored dots indicate the predicted activity level estimate within each season: Winter (choral), Spring (olive green), Summer (blue), Autumn (purple). Vertical lines depict the 95% confidence interval for the predicted activity level estimates for each season. Wider confidence intervals indicate greater variation among sites within each season. Asterisk (*) denotes significance to the level alpha <0.05.

**Table 2:** Pairwise seasonal differences in chacma baboon activity levels for each day period (Tukey’s post hoc comparisons following beta regression models).

|  | Dawn |  | Morning |  | Midday |  | Afternoon |  | Dusk |  | Night |  |
| --- | --- | --- | --- | --- | --- | --- | --- | --- | --- | --- | --- | --- |
|  | Coefficient estimate | P value | Coefficient estimate | P value | Coefficient estimate | P value | Coefficient estimate | P value | Coefficient estimate | P value | Coefficient estimate | P value |
| **Autumn - Winter** | 0.05 ± 0.24 | 0.9974 | 0.18 ± 0.13 | 0.4941 | -0.19 ± 0.18 | 0.6907 | -0.14 ± 0.19 | 0.8932 | -0.27 ± 0.24 | 0.6568 | -0.05 ± 0.27 | 0.9977 |
| **Autumn - Spring** | 1.04 ± 0.27 | 0.0006* | -0.04 ± 0.13 | 0.9913 | -0.23 ± 0.18 | 0.5793 | -0.20 ± 0.19 | 0.7276 | 0.32 ± 0.26 | 0.6075 | 0.68 ± 0.29 | 0.0782 |
| **Autum - Summer** | 2.24 ± 0.30 | < 0001* | -0.07 ± 0.13 | 0.9435 | 0.04 ± 0.20 | 0.9969 | 0.77 ± 0.24 | 0.0078* | 2.12 ± 0.30 | < 0001* | 1.86 ± 0.31 | < 0001* |
| **Spring - Summer** | 1.19 ± 0.30 | 0.0005* | -0.03 ± 0.13 | 0.9912 | 0.27 ± 0.19 | 0.4705 | 0.97 ± 0.23 | 0.0002* | 1.80 ± 0.30 | < 0001* | 1.18 ± 0.30 | 0.0006* |
| **Spring - Winter** | -1.00 ± 0.26 | 0.0006* | 0.22 ± 0.12 | 0.2883 | 0.04 ± 0.16 | 0.9963 | 0.07 ± 0.18 | 0.9827 | -0.59 ± 0.24 | 0.0628 | -0.73 ± 0.27 | 0.0324* |
| **Summer - Winter** | -2.19 ± 0.26 | < 0001* | 0.26 ± 0.13 | 0.2003 | -0.23 ± 0.18 | 0.5764 | -0.91 ± 0.23 | 0.0004* | -2.39 ± 0.29 | < 0001* | -1.91 ± 0.30 | < 0001* |

\* Denotes significance to the level $\alpha < 0.05$ .
Positive values indicate higher activity in the first season listed.

### Abiotic factors impact on the timing of chacma baboon’s activity (H3-4)

The observed per-site variations in diel activity level estimates of chacma baboons within each day period were primarily explained by variations in temperature (Fig. 5A, Fig. S4). Chacma baboons consistently showed lower activity levels in sites with higher temperatures throughout the day, although the effect was not significant during the morning (p=0.96; Fig. 5A) and at dawn (p=0.09; Fig. 5A). Additionally, precipitation emerged as a significant factor reducing activity level estimates at dawn (p<0.001) and at night (p=0.008; Fig. 5A). Finally, baboons were less active at midday when food availability was higher, as indicated by a significant negative effect of NDVI on activity levels (p=0.006; Fig. 5A, Fig. S4), with no notable impact observed during other periods of the day.

**Fig. 5.**
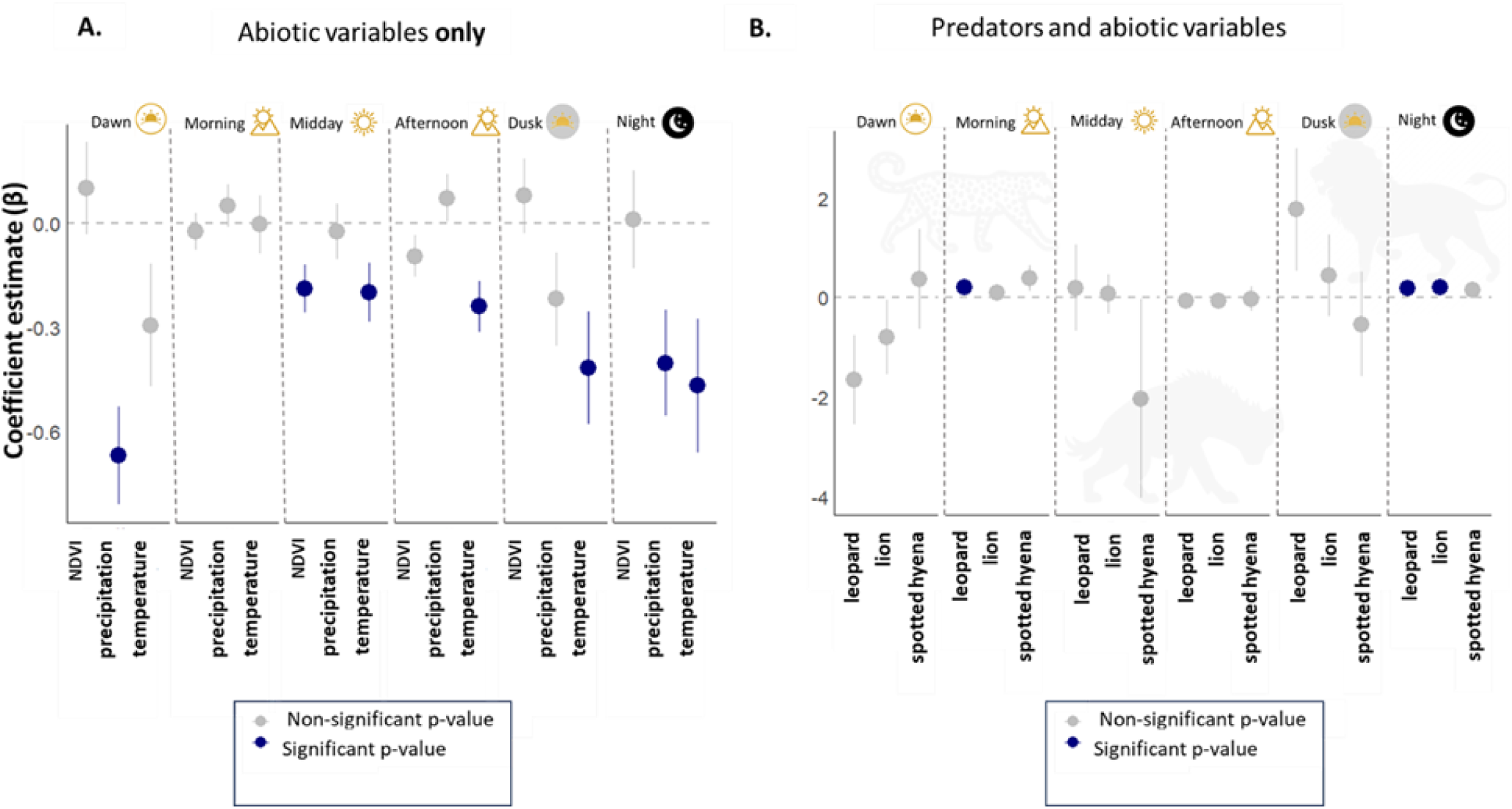
Drivers of variation in the total activity level estimates of chacma baboons in each day period: (**A).** Abiotic variables, namely temperature, precipitation and NDVI. (**B).** Biotic variables, namely lion, leopard and spotted hyena total activity level estimates (whilst controlling for temperature, precipitation and NDVI). The blue dots indicate mean coefficient estimate significantly different from 0 and the grey dots indicate non-significant mean coefficient estimates of beta regression at **α**=0.05. The bars represent 95% confidence intervals. Note the change of Y-axis scale between A and B.

### Predators’ influence on chacma baboon’s diel activity patterns (H5)

Predator temporal activity influenced chacma baboon activity levels in a pattern that varied by species and time of day, with varying degrees of temporal overlap across sites. Overall, chacma baboons showed a moderately low overlap with leopards (range = 0.14-0.55), with 16 out of 20 sites falling between 0.31±0.01 −0.55±0.01 (Fig. 6, Fig. S3). When considering day periods, leopards’ activity had the strongest effect on chacma baboon activity, showing a significant increase in chacma baboon activity at night (β = 0.20 ± 0.09, *p* = 0.03) and in the morning (β = 0.2±0.1, p = 0.05; Fig. 5B, fig. S4). Conversely, leopards showed an almost significant negative effect on chacma baboon activity at dawn (β = −1.7±0.9, p = 0.06; Fig. 5B).

**Fig. 6.**
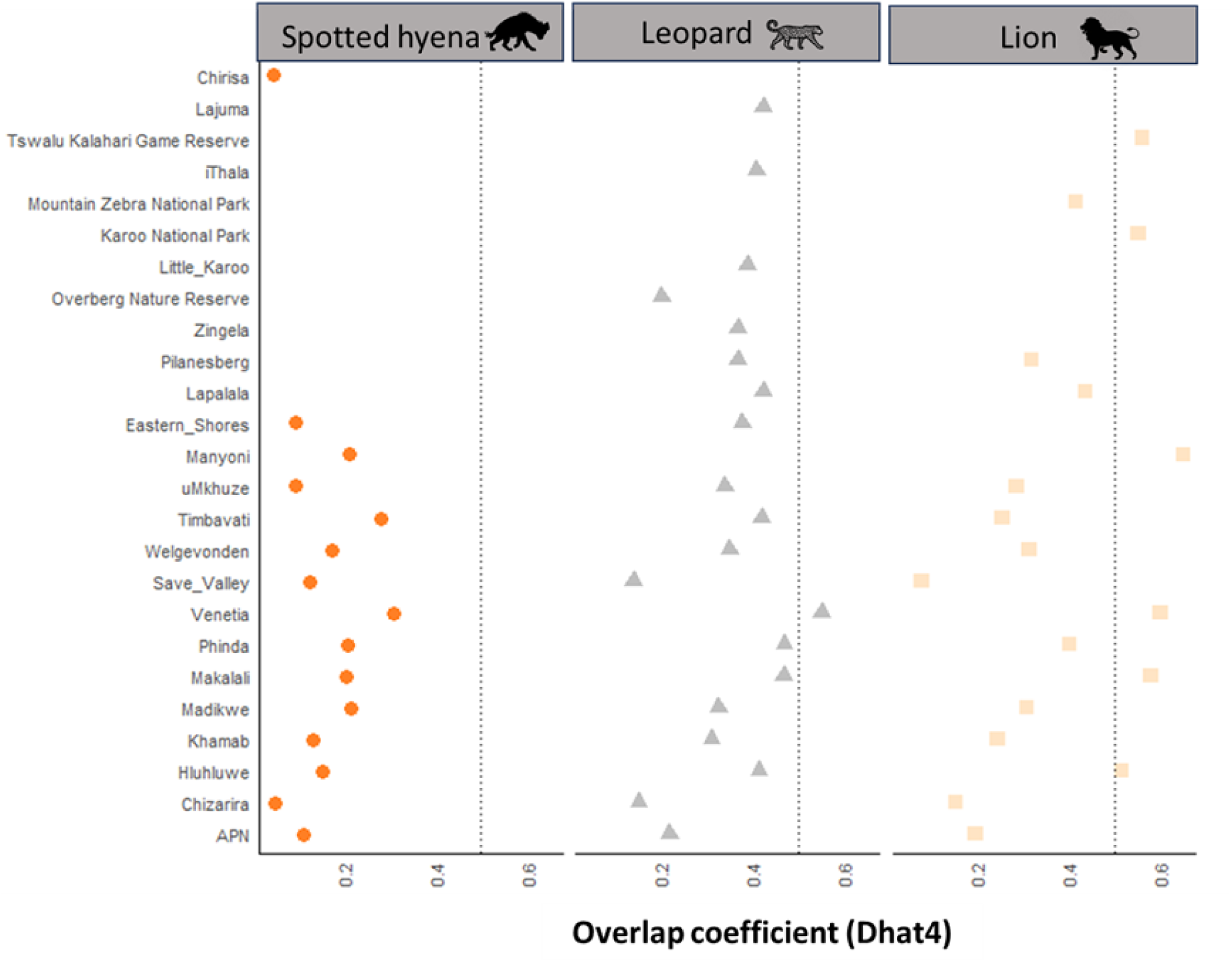
Overlap coefficient between chacma baboon and predator activity patterns in each site: spotted hyena (circles), leopard (triangles) and lion (squares). The y-axis lists the site names, while the x-axis shows the overlap coefficient (ranging from 0 = no overlap to 1 = complete overlap). A vertical dotted line at 0.5 to indicate moderate temporal overlap; values to the right of this line suggest relatively high temporal overlaps, while those to the left indicate a lower overlap. APN stands for Associate Private Nature Reserves.

Spotted hyenas had the lowest temporal overlap with chacma baboons (range=0.05-0.3 across all sites; Fig. 6, Fig. S3). Their effect was weak and non-significant across all tested day periods, although spotted hyena activity level estimates showed a marginally positive effect on chacma baboons’ activity patterns at night (β = 0.16 ± 0.07, *p* = 0.06), dawn (β = 0.37 ± 0.71 ± 1.00, *p* = 0.71) and morning (β = 0.39 ± 0.26, *p*= 0.13; Fig. 5B, Fig. S4).

Lions showed the highest temporal overlap with chacma baboons (range = 0.07-0.65 with a third of the sites having an overlap estimate of more than 0.52±0.01; Fig. 6, Fig. S3). Specifically, lion activity at night was associated with a significant increase in chacma baboons’ nocturnal activity (β = 0.21 ± 0.08, *p* = 0.01; Fig. 5B).

## Discussion

Time is a critical ecological factor influencing animal behavior and survival strategies. This study analyzed how environmental factors, temperature, food availability, and predation, shape the daily activity patterns of chacma baboons across diverse habitats. Utilizing seven years of camera trap data from 29 sites across six southern African biomes (Fig. 1), we identified landscape-level variations in baboon activity and their environmental drivers.

Throughout the day, chacma baboons exhibited a bimodal activity pattern, with peaks in the morning and afternoon, and reduced activity during midday, dusk, and night (Fig. 2, Fig. S2). This pattern, common among diurnal terrestrial mammals, reflects a thermoregulatory strategy to minimize heat stress and water loss (2,71). Similar bimodal activity rhythms have been observed in African elephants (*Loxodonta africana*), which rest during the hottest parts of the day to maintain homeostasis (72), and in red deer (*Cervus elaphus*), which adjust their activity to avoid thermal stress (20). Vervet monkeys (*Chlorocebus pygerythrus*) also modify their activity in response to temperature and predation risk (73). Notably, the extent of dawn and nocturnal activity varied among sites, particularly in environments with higher thermal or predation challenges, indicating behavioral flexibility in response to local conditions.

Consistent with our *geographical perspective* hypothesis (H1), total daily activity levels declined from southern to northern latitudes, with the highest levels recorded in the cooler, resource rich southern sites like Garden Route National Park (forest biome), and the lowest in hotter and more arid northern sites like Venetia (savanna biome; Fig. 2A). This variation reflects underlying environmental trade-offs on time-budget allocation. Within resource-rich, climatically stable environments, chacma baboons can sustain high activity levels over broader time windows due to lower thermoregulatory costs, more consistent precipitation and a high productivity year-round, supporting more consistent foraging returns (2,74). Conversely, in resource-poor or thermally stressful environments, such as the northern sub-Saharan Africa animals reduce activity or shift it to cooler periods to conserve energy and water. This mirrors patterns observed in Eurasian lynx (75), African lion (22) and other mammalian taxa that optimize energy use in response to environmental stress. Despite variation in total activity, wake-up (Fig. 3C) and sleep times (Fig. 3D) did not differ significantly across sites after adjusting for sunrise and sunset, indicating strong circadian entrainment. This suggests that while activity intensity varies, its timing is regulated by photoperiod (75). Slightly earlier wake-up and sleep times in some northern populations may reflect local predator or thermal pressures.

In partial contrast with our *seasonal perspective* hypothesis (H2), chacma baboons’ total daily activity level estimates did not differ significantly across seasons. However, the timing of activity varied seasonally as expected (Fig. 4). Specifically, activity during dawn, dusk, and night was significantly higher in winter and autumn compared to summer and spring, while morning and midday activity remained stable. This is potentially a compensatory strategy to meet energy demands in food-scarce months (39,76). Afternoon activity increased in winter and spring likely due to cooler temperatures; and decreased in summer to minimize heat stress (64), which aligns with our expectations. This seasonal redistribution of activity through the day, rather than a simple increase or decrease in total activity, reflects behavioral adjustments for energetic homeostasis and thermoregulations against fluctuating seasonal environmental pressures. Similar patterns are seen in diurnal non-human primate species like the Japanese macaques (*Macaca fuscata*) (77) and other mammal species such as the common brushtail possums (*Trichosurus vulpecula*) (78), that extend foraging into darker hours during short winter days to fulfill energetic demands. By reallocating activity rather than increasing effort, chacma baboons demonstrate adaptive responses that enhance ecological resilience and contribute to their success across heterogeneous landscapes.

Thermoregulation and foraging optimization are key physiological and behavioral processes influencing diel activity patterns(2,31). Supporting the *thermoregulatory constraints* hypothesis (H3), beta regression models identified temperature as the main abiotic factor shaping variation in activity across the day (Fig. 5A). Activity declined with increasing temperature, particularly at midday and afternoon which are the hottest periods of the day, while morning and dawn activity were not significantly impacted, likely due to cooler ambient conditions during those periods. As expected under the *food optimization* hypothesis (H4), NDVI was negatively correlated with mid-day activity, indicating that in food-rich areas, chacma baboons reduce foraging time during hot hours. Precipitation was also negatively associated with dawn and night activity, especially in drier regions, suggesting that baboons in resource-poor environments extend activity into low-light periods to meet energetic demands (Fig. 5A). Likewise, chacma baboons in the Cape Peninsula reduce mid-day activity when food is abundant to avoid prolonged foraging in heat-stressed hours (38). Similar strategies are observed in mammals inhabiting food scarce, hot and arid environments such as the African wild dogs (*L. pictus*), which increase activity during energetically costly hours to enhance their chances of locating prey (12,79), whereas the desert bighorn sheep (*Ovis canadensis nelsoni*) confine foraging to cooler dawn and dusk to avoid thermal stress, even at potential costs (18). These patterns illustrate how thermoregulation and food availability interact to shape when, and how much animals are active in a context-dependent manner.

Contrary to the *predation risk avoidance* hypothesis (H5), increased baboon activity at night and early morning coincided with the presence of their main predators: leopards, lions, and spotted hyenas. Leopard presence was associated with increased baboon activity at night and morning, while spotted hyenas had weaker, but similar effects (Fig. 5B). Unexpectedly, lions, often considered secondary baboon predators after leopards, had the strongest influence, significantly increasing nocturnal baboon activity and contributing to the highest predator-prey temporal overlap observed (Fig. 6, Fig. S3). These patterns suggest co-occurrence or reactive responses rather than strict temporal avoidance. These results challenge the classical model of predator avoidance and suggest more complex behavioral dynamics, such as vigilance, mobbing, or reorganization of sleeping sites in response to predator cues (41,42). Comparable reactive behaviors are seen in other systems. For instance, baboons in the Serengeti show group-level alarm and fleeing behavior following leopard encounters during the day (40,80). In ungulates, white-tailed deer (*Odocoileus virginianus*) adjust diel activity in response to wolves and hunters, often becoming more crepuscular or nocturnal depending on perceived risk (81), while African buffalo (*Syncerus caffer*) and zebra (*Equus quagga*) also adjust activity or group cohesion in response to lion predation pressure, but not always through strict temporal avoidance, highlighting the importance of trade-offs between safety, thermoregulation, and foraging needs (82).

The absence of clear temporal partitioning may reflect limited behavioral flexibility in diurnal species like chacma baboons (83). Rather than shifting core activity, they may rely on coexistence strategies such as spatial avoidance, group vigilance, and occasional nocturnal foraging (84). Temporal avoidance may also depend on predator hunting mode: leopards, as stealthy ambush hunters, may induce subtle timing shifts, while group-hunting lions and hyenas provoke more visible agitation, as seen in nocturnal activity (24). Temporal overlap was generally low with leopards and hyenas but more variable with lions, suggesting site- or season-specific dynamics.

This study utilizes predator activity levels as a proxy for predation pressure due to limitations in estimating predator abundance. While predator abundance is often used to assess predation intensity in mammalian systems, accurately determining it from camera trap data remains challenging (50,60). Reliable abundance estimates typically require intensive, targeted surveys and spatially explicit modelling approaches, which were beyond this study’s scope (8,85). Furthermore, seasonal variations in predator activity and prey vulnerability can influence these interactions. For instance, predators may increase hunting efforts during dry or resource-scarce seasons, prompting prey to adjust or extend their activity periods defensively. Future research incorporating seasonal predator movement data and group-level baboon behavior would help elucidate whether observed co-activity patterns are defensive, opportunistic, or constrained responses to shared diel activity.

## Conclusion

This study illustrates how environmental and ecological factors, specifically temperature, resource availability, and predator presence, influence diel activity patterns in chacma baboons across southern Africa. By analyzing data from 29 sites over multiple years, we provide a cross-population perspective on how baboons balance thermoregulation, foraging efficiency, and predator avoidance in their daily routines. Our results show that chacma baboons in climatically stable and resource-rich environments maintain higher activity levels, likely due to reduced energetic costs and expanded opportunities for behaviors such as social interaction and exploratory foraging. Conversely, in harsher or more seasonal environments, baboons exhibit compressed or shifted activity schedules, prioritizing essential foraging while minimizing thermal and predation risks. Importantly, this study highlights that diel activity strategies emerge from the interplay between ecological constraints and behavioral plasticity, a principle applicable across taxa. For instance, the predator species, particularly the presence of nocturnal or crepuscular predators, modulate baboon activity in complex ways, challenging simple models of temporal avoidance and suggesting context-dependent strategies. These findings underscore the value of viewing temporal niche adaptation not as a fixed species trait but as a flexible response to dynamic ecological pressures. By providing a mechanistic and spatially explicit understanding of diel activity variation, this study contributes to broader ecological theory on time-use strategies and adaptive behavior in variable environments.

## Acknowledgments

We express our gratitude to Panthera, the Cape Leopard Trust, and Snapshot Safari for providing access to camera trap data, along with their field technicians, students, and assistants for data collection and curation. Special thanks to Oriane Basso for assistance with downloading climatic data. We appreciate the support from the Nelson Mandela University Wildlife Ecology Lab Team. This work was supported by CNRS. The funders had no role in study design, data collection and analysis, decision to publish, or preparation of the manuscript. AI-assisted technologies were used for spelling and grammar check in this work.

## Funding

International Research Laboratory REHABS (LD)

CNRS, IRP RESONAB grant (EH)

Agence Nationale pour la Recherche ANR-22-CE92-0030-01 (EH)

National Research Foundation CSUR23031081837 (VR)

## Author contributions

Conceptualization: VR

Methodology: LD, VR, FP, LT, EH, MC

Investigation: HF, JF, AW, RG

Visualization: LD, VR

Funding acquisition: VR

Project administration: LD

Supervision: VR, FP, LT, EH

Writing – original draft: LD

Writing – review & editing: LD, VR, LT, FP, EH, HF, JF, MC, RG, AW

## Statement on inclusion

This study acknowledges the value of diverse perspectives and inclusivity by bringing together authors based on both countries where data was collected. Relevant literature from local scientists was actively cited. Where possible, preliminary findings have been presented to local researchers, conservation practitioners, and community members to invite feedback and foster dialogue.

## Data availability statement

The data used in this study will be available on Dryad repository at http://datadryad.org/share/juMSkCo_xBw7aAOdeyNiFLtwpnlFLJtmZbAAcC3l-QM. The raw camera-trap data employed in this study can be requested at Snapshot Safari (snapshotsafari.org), Cape Leopard Trust) and Panthera (panthera.org).

## Code availability

The code to analyze and reproduce this study is available in supplementary material.

