## Supplementary Information for "Climate and predation drive variation of diel activity patterns in chacma baboons (*Papio ursinus*) across southern Africa"

**The PDF file includes:**

Materials and Methods
Supplementary Text
Figs. S1 to S#
Tables S1 to S#
References

### Materials and Methods

#### Camera trap data

Snapshot Safari is dedicated to the comprehensive monitoring of wildlife across varied African ecosystems and provides extensive datasets for research (Pardo *et al.*, 2021). Panthera's southern Africa leopard monitoring program utilizes camera trapping to provide data on leopard population trends (Rogan *et al.*, 2022). Finally, Cape Leopard Trust's camera trap initiative is dedicated to the study and preservation of the cape leopards and other wildlife within the Western, Northern and Eastern Cape provinces of South Africa (Amin *et al.*, 2022). The primary objectives of these projects include gathering comprehensive data on animal presence, behavior, and population dynamics. These data are instrumental in informing conservation efforts (Pardo *et al.*, 2021), addressing human-wildlife conflicts (Puls *et al.*, 2021), conducting ecological studies (Amin *et al.*, 2022; Müller *et al.*, 2022), and facilitating biodiversity monitoring initiatives (Louw, Foden and Mann, 2020) For the Snapshot Safari project, infrared motion and heat-sensor cameras were systematically deployed following a standardized grid (for more details, refer to (Pardo *et al.*, 2021)). Each capture event consisted of a maximum of three images taken within five seconds of the trigger during daylight hours and one single image if the camera trap was triggered at night (Pardo *et al.*, 2021). Camera trap images were then meticulously classified and manually curated using both citizen science (Zooniverse; https://www.zooniverse.org) and trained laboratory assistance using Traptagger (https://wildeyeconservation.org/trap-tagger-about/). In the case of the Cape Leopard Trust and Panthera projects, paired cameras were positioned facing each other at each station. Cape Leopard Trust cameras were set to take a burst of three images per trigger during the day, and one image at night. For the current study, in cases where the animal was detected by both cameras, records from only one camera were retained for each station to avoid duplicates in areas with high animal detectability, such as roads, drainage lines, and animal paths (Louw, Foden and Mann, 2020; Amin *et al.*, 2022). The Cape Leopard Trust data was classified using WildID (https://www.wildid.app/), a machine learning program that automatically identifies and labels animal species from images, and then user verified. Images from the Panthera surveys were classified into species both by research team members and a machine learning algorithm, using Panthera's Integrated Data Software, PantheraIDS, a custom-built camera trap data processing package, within the R Statistical Environment (R Core Team, 2024).

**Fig. S1.**

**Filtering and dataset processing.** Each box in the diagram represents the number of sites and indicates the number of remaining images for each project (Panthera, Snapshot Safari, and Cape Leopard Trust**).** Key filtering options are also indicated in the ovals. The final steps show the analyses performed using the different datasets.

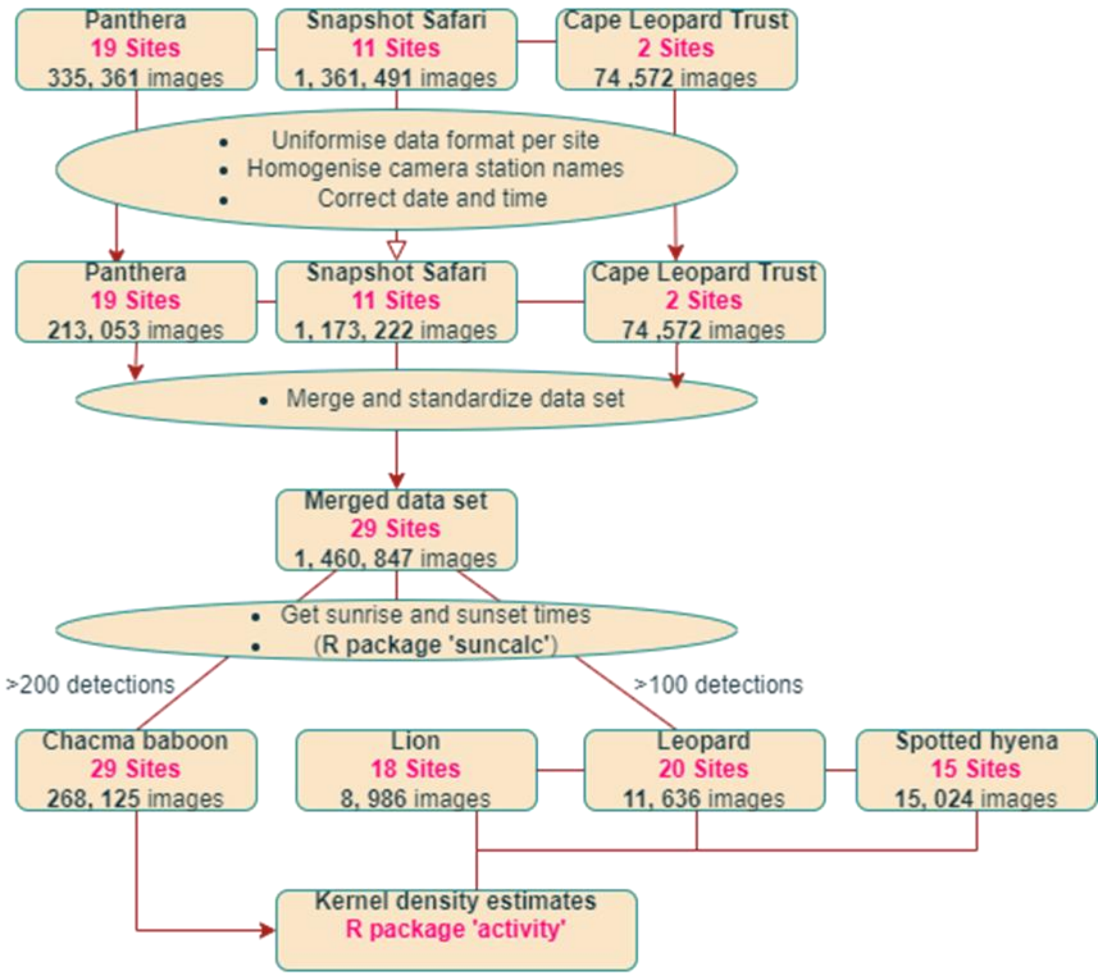

**Fig. S2.**

**Chacma baboons' activity density patterns in each of the 29 study sites.** The Y-axis represents the density of activity, and the X-axis represents the hours throughout the day (solar time). Each day is divided into six periods: dawn and dusk (grey), morning and afternoon (yellow), midday (orange), night (black).

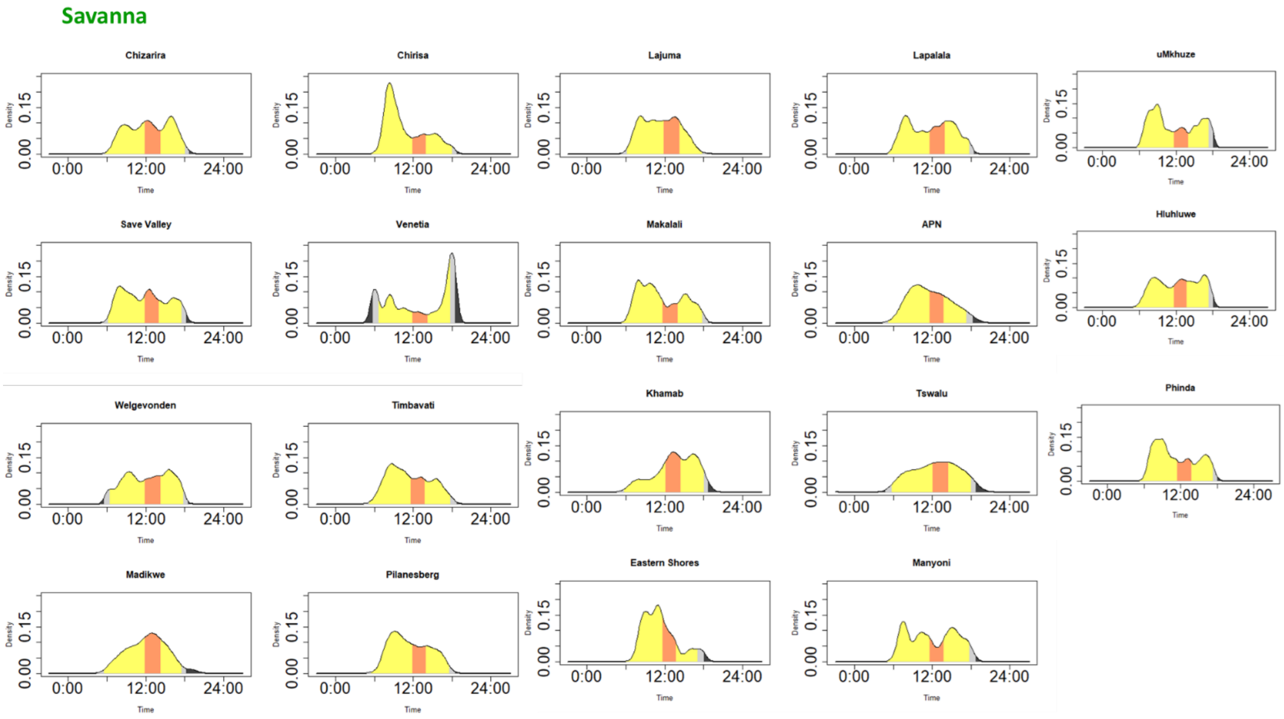

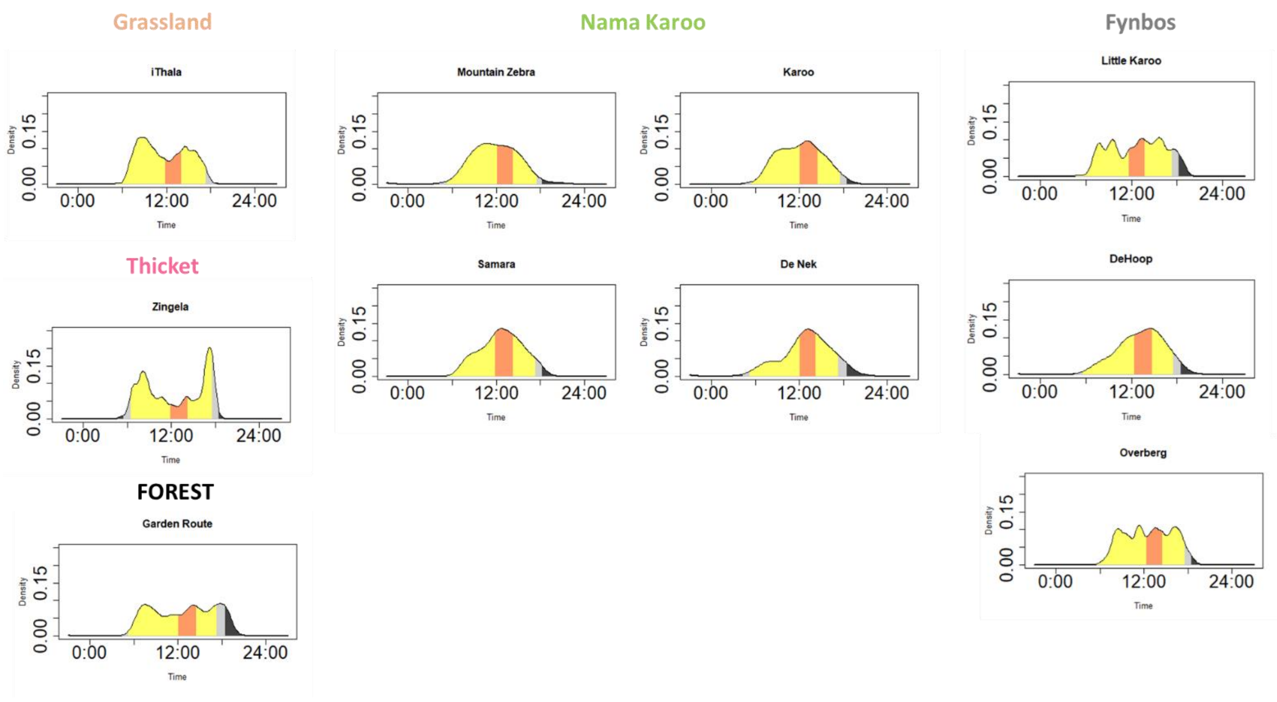
Grassland
iThala
Nama Karoo
Mountain Zebra
Karoo
Samara
De Nek
Fynbos
Little Karoo
DeHoop
Overberg
Thicket
Zingela
FOREST
Garden Route
Density
0.00
0.15
0:00
12:00
24:00
Time

**Fig. S3.**

**Overlap between chacma baboons and predators' activity density patterns in each of the 29 study sites.** The Y-axis represents the density of activity, and the X-axis represents the hours throughout the day (solar time, see text). Each line represents a species: chacma baboon (brown), leopard (grey), lion (light peach), spotted hyena (orange).

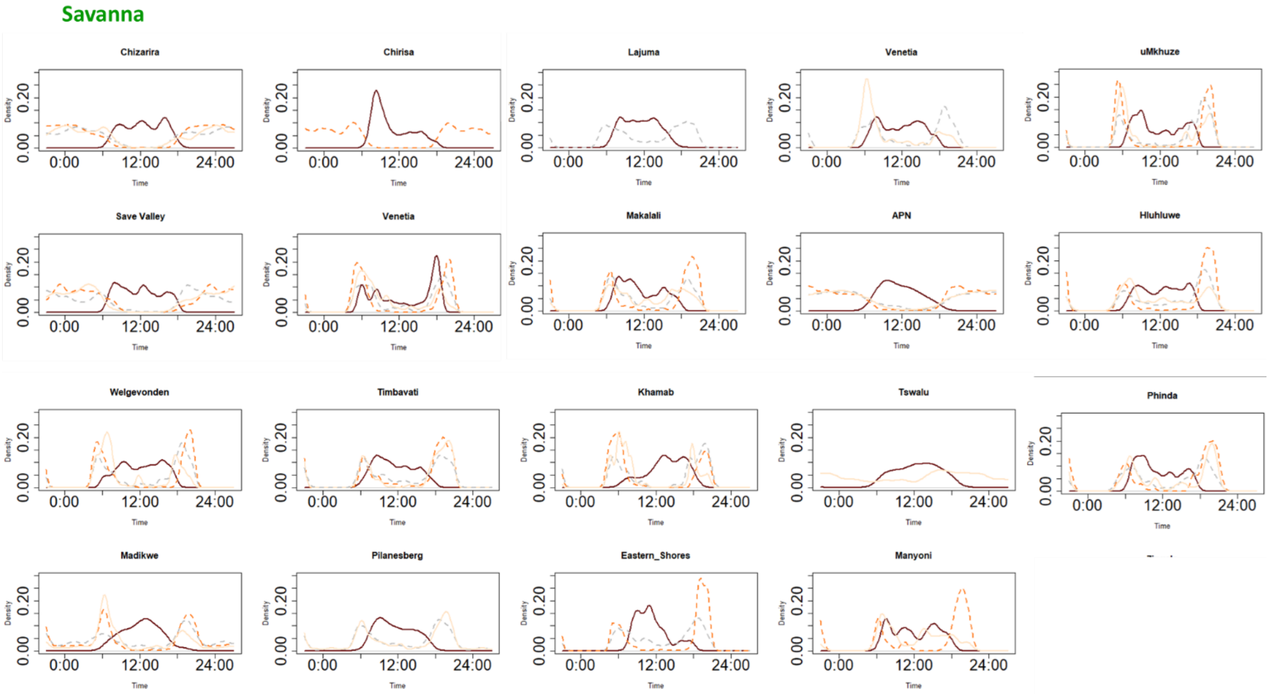

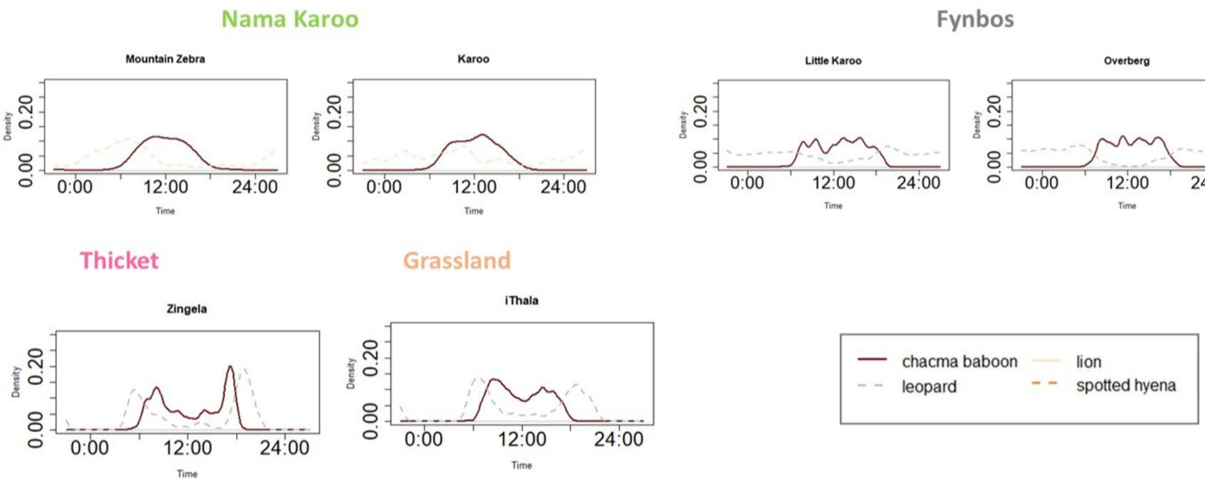
Nama Karoo
Fynbos
Mountain Zebra
Karoo
Little Karoo
Overberg
Thicket
Grassland
Zingela
iThala
Density
0.00
0.20
0:00
12:00
24:00
Time
chacma baboon
lion
leopard
spotted hyena

**Fig. S4.**

**Extended drivers of variation in the total activity level estimates of chacma baboons in each day period.** The Y-axis represents activity level estimates of chacma baboons, and the X-axis represents scaled values of each abiotic driver (temperature, precipitation, NDVI) and activity level estimates of predators (lion, leopard, spotted hyena). Circles represent activity level estimates at dawn (maroon), morning (green), midday (purple), afternoon (blue), dusk (olive green), and night (brown). Lines represent predictions from the Beta regression models along a range of activity level estimates. The shaded area represents 95% confidence intervals at $\alpha$=0.05.

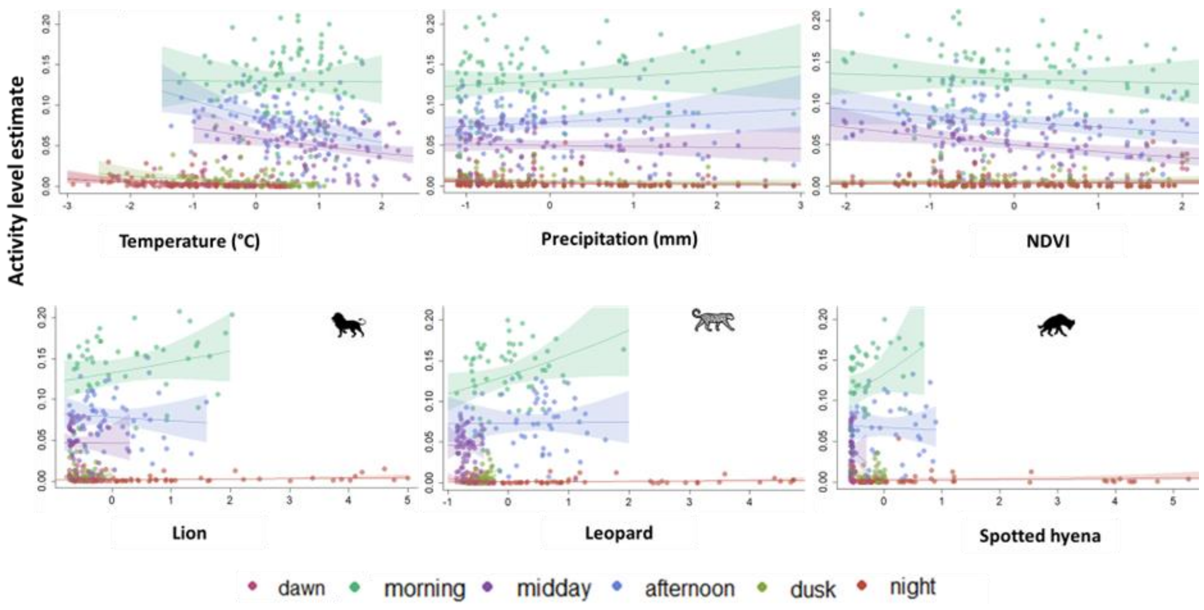

**Table S1.**

**Description of the 29 study sites used for camera trap data collection.** For each site, we provide the country, biome classification, mean summer and winter temperatures (in °C), mean annual rainfall (in mm), main predators of chacma baboons, dominant plant species, reserve size (in hectares), rainfall seasonality (summer, winter, or year-round), and type of land management (e.g., privately owned, state or provincial government, national government). Predator presence is based on confirmed detections or known historical occurrences at each site. Plant species listed reflect the dominant vegetation type observed within the main baboon habitat areas. NA. Not available.

| Number ID | Site | Country | Biome | Mean summer temperature (°C) | Mean winter temperature (°C) | Mean annual rainfall (mm) | Mean annual NDVI | Key baboon predators | Dominant plant species | Size (ha) | Rainfall seasonality | Management |
|---|---|---|---|---|---|---|---|---|---|---|---|---|
| 1 | Venetia Limpopo Nature Reserve | South Africa | Savanna | 26 | 16 | 366 | 0.34 | spotted hyena, lion, leopard | *Colophospermum mopane* | 39000 | Summer | Privately owned |
| 2 | Lajuma Research Centre | South Africa | Savanna | 24 | 11 | 724 | 0.59 | spotted hyena, leopard | | 430 | Summer | Privately owned |

| | | | | | | | | | | | | |
|---|---|---|---|---|---|---|---|---|---|---|---|---|
| 3 | Welgevonden Game Reserve | South Africa | Savanna | 23 | 13 | 665 | 0.46 | spotted hyena, lion, leopard | | 37500 | Summer | Privately owned |
| 4 | Timbavati Private Nature Reserve | South Africa | Savanna | 28 | 19 | 550 | 0.39 | spotted hyena, lion, leopard | *Terminalia sericea, Colophospermum mopane, Combretum. Spp* | 53392 | Summer | Privately owned |
| 5 | Makalali Game Reserve | South Africa | Savanna | 32 | 18 | 450 | 0.45 | spotted hyena, lion, leopard | | 26000 | Summer | Privately owned |
| 6 | Lapalala Wilderness | South Africa | Savanna | 22 | 12 | 500 | 0.44 | spotted hyena, lion, leopard | | 48000 | Summer | Privately owned |
| 7 | Associate Private Nature Reserves | South Africa | Savanna | 26 | 19 | 550 | 0.37 | spotted hyena, lion, leopard | | 182500 | Summer | Privately owned |
| 8 | Pilanesberg National Park | South Africa | Savanna | 29 | 13 | 630 | 0.44 | spotted hyena, lion, leopard | | 57200 | Summer | Provincial government |

| | | | | | | | | | | | | |
|---|---|---|---|---|---|---|---|---|---|---|---|---|
| 9 | Madikwe Game Reserve | South Africa | Savanna | 30 | 13 | 750 | 0.40 | spotted hyena, lion, leopard | | 75000 | Summer | Provincial government |
| 10 | Khamab Kalahari Reserve | South Africa | Savanna | 26 | 11 | 330 | 0.28 | spotted hyena, lion, leopard | | 95538 | Summer | Privately owned |
| 11 | Tswalu Kalahari Reserve | South Africa | Savanna | 34 | 10 | 253 | 0.28 | spotted hyena, lion, leopard | *Vachellia erioloba, Boscia albitrunca and Terminalia sericea* | 102000 | Summer | Privately owned |
| 12 | uMkhuze Game Reserve | South Africa | Savanna | 32 | 18 | 1200 | 0.57 | spotted hyena, lion, leopard | | 40000 | Summer | Provincial government |
| 13 | Phinda Private Game Reserve | South Africa | Savanna | 30 | 18 | 1000 | 0.57 | spotted hyena, lion, leopard | | 28555 | Summer | Privately owned |
| 14 | Manyoni Private Game reserve | South Africa | Savanna | 26 | 16 | 580 | 0.60 | spotted hyena, lion, leopard | | 21700 | Summer | Privately owned |

| | | | | | | | | | | | | |
|---|---|---|---|---|---|---|---|---|---|---|---|---|
| 15 | Ithala Game Reserve | South Africa | Grassland | 24 | 20 | 791 | 0.55 | spotted hyena, lion, leopard | | 29653 | Summer | Provincial government |
| 16 | Hluhluwe Game Reserve | South Africa | Savanna | 25 | 17 | 620 | 0.61 | spotted hyena, lion, leopard | | 95000 | Summer | Provincial government |
| 17 | Eastern Shores | South Africa | Savanna | 30 | 20 | 1129 | 0.66 | spotted hyena, leopard | | 264000 | Summer | Provincial government |
| 18 | Zingela Nature Reserve | South Africa | Thicket | 36 | 12 | 421 | 0.36 | spotted hyena, lion, leopard | | 27000 | Summer | Provincial government |
| 19 | Samara Private Game Reserve | South Africa | Nama Karoo | 22 | 13 | 336 | 0.36 | lion | *Acacia karroo, Themeda tiandra* | 28000 | Summer | Privately owned |
| 20 | DeNek Farm | South Africa | Nama Karoo | 22 | 13 | 336 | 0.31 | absent | | unknown | Summer | Privately owned |

| | | | | | | | | | | | | |
|---|---|---|---|---|---|---|---|---|---|---|---|---|
| 21 | Mountain Zebra National Park | South Africa | Nama Karoo | 29 | 15 | 400 | 0.31 | lion | | 28412 | Summer | National government |
| 22 | Karoo National Park | South Africa | Nama Karoo | 27 | 11 | 220 | 0.13 | spotted hyena, lion | | 76990 | Summer | National government |
| 23 | Overberg Nature Reserve | South Africa | Fynbos | 22 | 13 | 600 | 0.42 | leopard | *Proteas, Ericas and Restios* | | Winter | Provincial government |
| 24 | Little Karoo | South Africa | Nama Karoo | 21 | 14 | 220 | 0.33 | leopard | | 62000 | Summer | Privately owned |
| 25 | Garden Route National Park | South Africa | Forest | 15 | 18 | 1000 | 0.74 | leopard | | 130000 | All year | National government |
| 26 | DeHoop Nature Reserve | South Africa | Fynbos | 21 | 14 | 455 | 0.53 | absent | *Proteas, Ericas and Restios* | 36000 | Winter | Provincial government |
| 27 | Save Valley Conservancy | Zimbabwe | Savanna | 33 | 18 | 450 | 0.40 | spotted hyena, lion, leopard | | 344200 | Summer | Privately owned |

| | | | | | | | | | | | | |
|---|---|---|---|---|---|---|---|---|---|---|---|---|
| 28 | Chirisa Safari Area | Zimbabwe | Savanna | 28 | 13 | 600 | 0.37 | spotted hyena, lion, leopard | *Colophospermum mopane, Acacia spp, Combretum spp* | 171300 | Summer | State owned |
| 29 | Chizaria National Park | Zimbabwe | Savanna | 24 | 18 | 800 | 0.37 | spotted hyena, lion, leopard | *Colophospermum mopane, Terminalia spp* | 200000 | Summer | State owned |

**Table S2.**

**Summary of camera trap deployment location and capture data for chacma baboons (*Papio ursinus*) and their primary predators (spotted hyena, leopard, and lion) across 29 study sites in South Africa and Zimbabwe.** For each site are indicated the site name, source project, country, biome, number of camera traps deployed (CTs), start and end dates of monitoring, total number of trap days, and total number of capture events. "Unfiltered" and "Filtered" refer to raw and manually verified capture events, respectively. Predator and baboon capture columns correspond to capture events after filtration.

| Number ID | Site name | Source project | Country | Biome | CTs | Start date | End date | Trap days | Unfiltered | Filtered | Chacma baboon | Spotted hyena | Leopard | Lion |
|---|---|---|---|---|---|---|---|---|---|---|---|---|---|---|
| 1 | Venetia Limpopo Nature Reserve | Panthera | South Africa | Savanna | 40 | 02/07/2016 | 06/07/2022 | 197 | 11,531 | 6,002 | 4,342 | 146 | 409 | 471 |
| 2 | Lajuma Nature Reserve | Panthera | South Africa | Savanna | 40 | 08/03/2016 | 19/05/2022 | 412 | 66,489 | 39,093 | 37,982 | 22 | 853 | 0 |
| 3 | Welgevonden Game Reserve | Panthera | South Africa | Savanna | 40 | 06/04/2016 | 17/07/2022 | 348 | 25,333 | 14,284 | 11,975 | 359 | 698 | 729 |
| 4 | Timbavati Private Nature Reserve | Panthera | South Africa | Savanna | 40 | 01/09/2016 | 08/11/2022 | 291 | 15,569 | 6,644 | 2,669 | 3173 | 571 | 247 |
| 5 | Makalali Game Reserve | Panthera | South Africa | Savanna | 40 | 03/09/2016 | 31/12/2022 | 356 | 19,119 | 9,602 | 5,874 | 1,520 | 804 | 1,191 |
| 6 | Lapalala Wilderness | Panthera | South Africa | Savanna | 40 | 29/10/2016 | 13/09/2022 | 323 | 13,446 | 11,336 | 9,435 | 51 | 532 | 308 |

| | | | | | | | | | | | | | | |
|---|---|---|---|---|---|---|---|---|---|---|---|---|---|---|
| 7 | Associate Private Nature Reserves | Snapshot Safari | South Africa | Savanna | 45 | 28/06/2017 | 26/11/2019 | 880 | 51,401 | 47,810 | 476 | 1859 | 168 | 246 |
| 8 | Pilanesberg National Park | Panthera | South Africa | Savanna | 40 | 13/03/2016 | 14/03/2022 | 286 | 16,398 | 6,763 | 1,394 | 38 | 567 | 1102 |
| 8 | Pilanesberg National Park | Snapshot Safari | South Africa | Savanna | 39 | 07/10/2017 | 26/02/2021 | 1681 | 31,253 | 31,237 | 236 | 14 | 67 | 245 |
| 9 | Madikwe Game Reserve | Panthera | South Africa | Savanna | 40 | 01/11/2016 | 30/10/2022 | 338 | 11,714 | 6,764 | 3,256 | 1077 | 141 | 412 |
| 9 | Madikwe Game Reserve | Snapshot Safari | South Africa | Savanna | 53 | 11/06/2018 | 23/02/2021 | 1023 | 141,363 | 123,458 | 1,322 | 384 | 100 | 145 |
| 10 | Khamab Kalahari Reserve | Panthera | South Africa | Savanna | 40 | 01/09/2016 | 30/08/2022 | 1119 | 7,702 | 4,726 | 1,505 | 271 | 335 | 1,303 |
| 11 | Tswalu Kalahari Reserve | Snapshot Safari | South Africa | Savanna | 30 | 18/11/2018 | 16/09/2022 | 1410 | 21,3052 | 107,706 | 769 | 16 | 16 | 175 |
| 12 | uMkhuze Game Reserve | Panthera | South Africa | Savanna | 40 | 27/05/2016 | 19/08/2022 | 337 | 18,002 | 11,586 | 9,568 | 541 | 1177 | 297 |
| 13 | Phinda Private Game Reserve | Panthera | South Africa | Savanna | 42 | 24/06/2016 | 16/12/2019 | 175 | 10,972 | 7,162 | 6,167 | 301 | 368 | 327 |
| 14 | Little Karoo | Cape Leopard Trust | South Africa | Nama Karoo | 61 | 22/07/2022 | 29/12/2022 | 160 | 23,589 | 23,589 | 22,994 | 0 | 600 | 0 |
| 14 | Manyoni Private Game Reserve | Panthera | South Africa | Savanna | 37 | 15/02/2017 | 15/12/2022 | 148 | 2,743 | 1,721 | 1,198 | 116 | 63 | 312 |
| 15 | Ithala Game Reserve | Panthera | South Africa | Grassland | 50 | 23/07/2016 | 02/03/2022 | 350 | 51,721 | 42,373 | 40,678 | 21 | 1,354 | 2 |
| 16 | Hluhluwe Game Reserve | Panthera | South Africa | Savanna | 46 | 01/04/2016 | 17/10/2022 | 492 | 22,967 | 14,232 | 12,194 | 1,197 | 356 | 503 |
| 17 | Eastern Shores | Panthera | South Africa | Savanna | 41 | 16/09/2016 | 24/06/2022 | 350 | 12,282 | 5,590 | 1,740 | 2,639 | 1,210 | 0 |
| 18 | Zingela Nature Reserve | Panthera | South Africa | Thicket | 41 | 02/05/2016 | 10/05/2022 | 285 | 16,910 | 12,712 | 11,676 | 13 | 236 | 1 |

| | | | | | | | | | | | | | |
|---|---|---|---|---|---|---|---|---|---|---|---|---|---|
| 19 | Samara Private Game Reserve | Snapshot Safaris | South Africa | Nama Karoo | 20 | 28/05/2016 | 09/02/2020 | 313 | 15,401 | 15,392 | 2,263 | 0 | 0 | 6 |
| 20 | DeNek Farm | Snapshot Safari | South Africa | Nama Karoo | 9 | 24/06/2019 | 14/09/2020 | 384 | 4,673 | 4,673 | 242 | 0 | 0 | 0 |
| 21 | Mountain Zebra National Park | Snapshot Safari | South Africa | Nama Karoo | 19 | 10/04/2017 | 01/04/2021 | 1470 | 112,789 | 112,706 | 1,143 | 0 | 0 | 103 |
| 22 | Karoo National Park | Snapshot Safari | South Africa | Nama Karoo | 25 | 04/10/2017 | 19/10/2020 | 1035 | 66,286 | 13,294 | 12,710 | 1 | 0 | 348 |
| 23 | Overberg | Cape Leopard Trust | South Africa | Fynbos | 77 | 02/08/2021 | 10/02/2022 | 127 | 50,983 | 50,983 | 50,417 | 0 | 547 | 0 |
| 23 | Overberg Nature Reserve | Snapshot Safari | South Africa | Fynbos | 18 | 05/09/2018 | 05/07/2021 | 910 | 22,328 | 17,628 | 629 | 0 | 11 | 0 |
| 25 | Garden Route National Park | Snapshot Safari | South Africa | Forest | 73 | 01/01/2018 | 20/05/2022 | 491 | 685,940 | 682,313 | 3,161 | 0 | 64 | 0 |
| 26 | DeHoop Nature Reserve | Snapshot Safari | South Africa | Fynbos | 38 | 08/09/2018 | 09/11/2019 | 438 | 17,005 | 17,005 | 601 | 0 | 0 | 0 |
| 27 | Save Valley Conservancy | Panthera | Zimbabwe | Savanna | 40 | 14/07/2020 | 10/08/2022 | 239 | 8,944 | 8,944 | 7,317 | 680 | 341 | 239 |
| 28 | Chirisa Safari Area | Panthera | Zimbabwe | Savanna | 34 | 04/07/2019 | 16/11/2022 | 124 | 1,525 | 1,525 | 1,071 | 366 | 70 | 4 |
| 29 | Chizaria National Park | Panthera | Zimbabwe | Savanna | 38 | 29/06/2019 | 26/10/2022 | 131 | 1,994 | 1,994 | 1,121 | 395 | 191 | 283 |
| | | | | | | | **Total** | **16,623** | **1,771,424** | **1,460,847** | **268,125** | **15,200** | **11,849** | **8,999** |
